# Variability amongst maize genotypes treated with neonicotinoid and stored

**DOI:** 10.64898/2026.03.04.709654

**Authors:** Venicius Urbano Vilela Reis, Giulyana Isabele Silva Tavares, Marina Silva Resende Pereira, Suemar Alexandre Gonçalves Avelar, Marcelo Angelo Cirillo, Genaina Aparecida de Souza, Everson Reis Carvalho

## Abstract

Neonicotinoid insecticides used in seed treatment present phytotoxic potential that may accelerate seed deterioration during storage; however, how this effect interacts with genotype remains poorly understood. We evaluated the physiological quality of five maize genotypes sourced from the Maize Breeding Programme of the Federal University of Lavras, comprising three inbred lines (L44, L91 and L64) and two half-sib hybrids (H44 and H91), treated with a neonicotinoid-based insecticide formulation (thiamethoxam and cyantraniliprole) and stored for up to nine months at 25°C. Physiological quality was assessed through germination on rolled paper + vermiculite, cold test, primary root length, a phytotoxicity index, projection pursuit multivariate analysis, and scanning electron microscopy of pericarp and aleurone layer. Insecticide treatment reduced germination and increased phytotoxicity indices, with inbred line L44 showing the most severe response, reaching phytotoxicity values up to 15.89 percentage points above hybrid H91 at six months of storage. Scanning electron microscopy revealed that L44 presented a thinner pericarp and pronounced aleurone layer disorganisation following treatment, whilst L91 remained structurally preserved. Tolerance to post-treatment storage is highly genotype-dependent, and pericarp thickness may represent a useful morphological marker for selecting tolerant genotypes in maize breeding programmes.

## Introduction

With the continuous growth of the world population, the demand for food, feed, fibre and bioenergy has increased significantly, being met primarily by crops such as maize, rice and wheat, which are highly susceptible to biotic and abiotic stresses (Loogen *et al*., 2021).

In this context, the use of high-quality maize (*Zea mays* L.) seed is essential to ensure adequate early plant establishment, thereby favouring high yields (Reis *et al*., 2022; Reed *et al*., 2022). Research into methods that ensure seedling protection against pest and pathogen attack during the initial growth stage, referred to as seed treatment (ST), is therefore of paramount importance (Brustolin *et al*., 2017).

This technique involves the application of plant protection products, dyes, film coatings or additives directly onto the seed coat, protecting the seeds and allowing the full expression of their genetic potential in the field, whilst also reducing environmental impact compared with spraying (Reis *et al*., 2026).

Notwithstanding its operational and environmental benefits, this practice is subject to limitations. Among the principal challenges is the potential phytotoxicity of certain plant protection products, which may accelerate seed deterioration during storage. Such effects may result in damage to morpho-anatomical and histochemical structures, disruption of physiological and biochemical processes, and alterations in the homeostasis of reactive oxygen species (ROS) in seeds (Fatma *et al*., 2018; Rodrigues *et al*., 2019; Oliveira *et al*., 2020).

This effect is directly dependent on the active ingredient used and on the environmental conditions during storage, representing a significant obstacle to the maintenance of the physiological quality of treated and stored seeds (Moraes *et al*., 2022; Reis *et al*., 2023; Medeiros *et al*., 2026).

With the growing adoption of ST in the market, there has been an increasing use of insecticidal molecules, such as neonicotinoids, which are important tools in integrated pest management owing to their efficacy. However, the use of these molecules warrants attention, as they present a greater potential for phytotoxicity when seeds are stored for extended periods (Tamindžić *et al*., 2016; Medeiros *et al*., 2026; Reis *et al*., 2026).

Despite the advanced technologies applied to maize seed production and processing, significant knowledge gaps remain regarding how to guarantee and maintain maximum seed quality. The maintenance of seed physiological quality is considerably influenced by genotype, particularly under stress conditions such as those caused by treatment and storage with neonicotinoid insecticides (Tamindžić *et al*., 2016; Santos *et al*., 2017).

Studies relating genotype to initial seed physiological quality, and to the impacts of deterioration that intensify following storage, are therefore relevant (Ghosh *et al*., 2020; Olasoji and Ogunniyan, 2020; Yang *et al*., 2025). Each genotype confers unique physical, morphological, physiological, biochemical and metabolic characteristics to seeds (Saatkamp *et al*., 2019), directly affecting essential functions such as viability, longevity and deterioration control (Arif *et al*., 2022).

This inter-genotypic variability significantly affects responses to both biotic and abiotic stresses, demanding innovative solutions. The current body of literature lacks studies that comprehensively evaluate how different plant protection products and genotypes interact and affect seed physiological quality following storage, thereby assessing the variability of this trait.

Seed treatment with neonicotinoid insecticides in stored seeds may interact with the genetic constitution of the seeds during storage. The present study aimed to evaluate the variability in physiological quality and phytotoxicity response of maize genotypes subjected to neonicotinoid-based seed treatment and stored for up to nine months, representing a pioneering and innovative contribution with findings potentially essential to maximising the productive capacity of the maize crop.

## Material and methods

A completely randomised design was employed in a 5 × 2 × 4 factorial arrangement (genotypes, seed treatments and storage periods, respectively).

### Biological materials

Three inbred lines and two half-sib families of maize seeds were selected for this experiment, sourced from the germplasm belonging to the Maize Breeding Programme of the Federal University of Lavras (MBP/UFLA).

Of the three inbred lines, two were selected as female parents (L44 and L91) and one as the male parent (L64) for the production of the half-sib families (hybrids H44 and H91), based on contrasting characteristics regarding seed physiological quality, as identified in previous research on genotype tolerance to water deficit.

The half-sib families and inbred lines were produced during the 2021/2022 growing season by MBP/UFLA. Following manual harvesting, ears were dried in a dryer to 12% moisture content, after which seeds were shelled, cleaned and classified using round-hole sieves of 18 and 19 mm.

### Seed treatment and storage conditions

All seeds were initially treated with a standard slurry (control), comprising a fungicide mixture (azoxystrobin + thiabendazole + fludioxonil + metalaxyl-M) and a polymer (Disco AG Red L-450®, density 1.05–1.15 g mL^⁻1^, viscosity 300–1000 cPs at 25°C, pH 6–8) (Table 1).

**Table 1.**
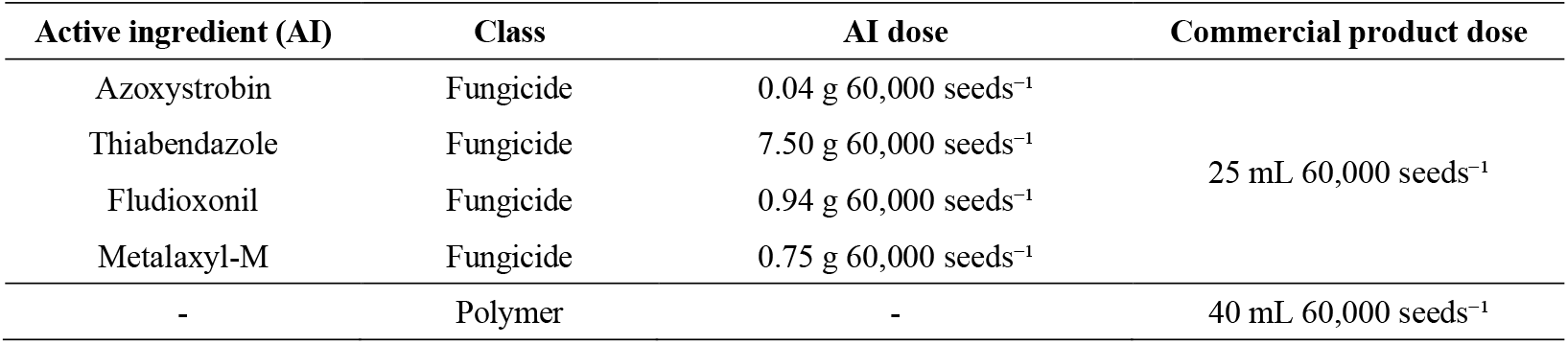
Composition of the standard slurry (control) used in the treatment of maize seeds.

| Active ingredient (AI) | Class | AI dose | Commercial product dose |
| --- | --- | --- | --- |
| Azoxystrobin | Fungicide | 0.04 g 60,000 seeds<sup>-1</sup> | 25 mL 60,000 seeds<sup>-1</sup> |
| Thiabendazole | Fungicide | 7.50 g 60,000 seeds<sup>-1</sup> |  |
| Fludioxonil | Fungicide | 0.94 g 60,000 seeds<sup>-1</sup> |  |
| Metalaxyl-M | Fungicide | 0.75 g 60,000 seeds<sup>-1</sup> |  |
| - | Polymer | - | 40 mL 60,000 seeds<sup>-1</sup> |

Insecticides were subsequently added to complete the Fortenza® Duo (FD) slurry, with the active ingredients (AI) thiamethoxam (a systemic and ingestion insecticide belonging to the neonicotinoid chemical group) and cyantraniliprole (a systemic contact and ingestion insecticide belonging to the anthranilic diamide chemical group), an insecticide considered unlikely to cause acute damage to seeds (Table 2).

**Table 2.** Insecticide products and doses used in the treatment of maize seeds.

| Active ingredient (AI) | AI dose | Commercial product dose |
| --- | --- | --- |
| Thiamethoxam | 3.60 g 60,000 seeds<sup>-1</sup> | 60 mL 60,000 seeds<sup>-1</sup> |
| Cyantraniliprole | 4.20 g 60,000 seeds<sup>-1</sup> | 70 mL 60,000 seeds<sup>-1</sup> |

Seeds were treated in 1.2 kg batches using a Momesso Arktos Laboratório L2K BM equipment, whose frequency inverter was calibrated to 15 Hz to simulate industrial treatment conditions. The complete application and homogenisation process for each batch lasted 20 seconds.

Following an air-drying period of three days at ambient temperature, the treated seeds were placed into storage. Each 1.2 kg batch, previously allocated to its respective evaluation period, was packed in multi-layer paper bags. Storage was carried out in BOD (Biochemical Oxygen Demand) chambers maintained at a constant temperature of 25 ± 1°C throughout the experimental period.

### Physical and physiological quality evaluations

Assessments of seed physiological quality, as described below, were conducted at 0, 2, 6 and 9 months after storage.

- Moisture content: was determined according to the methodology prescribed in the Rules for Seed Testing, RAS (Brasil, 2025), with results expressed as a percentage.
- Germination on rolled paper plus vermiculite (RP+V): the test was conducted with four replicates of 50 seeds. Seeds were sown on a sheet of germination paper moistened with water equivalent to three times the paper’s weight, after which 100 mL of moist vermiculite (1:1 w/v ratio) was distributed uniformly over the seeds. Seeds were then covered with a second sheet of germination paper, rolled, and maintained at 25 ± 2°C in a germination chamber. Seedling assessments were performed four and seven days after sowing, based on the percentage of normal seedlings (Rocha *et al*., 2023).
- Cold test (CT): the test was conducted with four replicates of 50 seeds per treatment. Seeds were sown in a sand and soil substrate at a 2:1 (v/v) ratio, moistened to 60% of its water-holding capacity. Following sowing, plastic trays (51 × 30 × 9 cm) containing the substrate were kept in a cold chamber at 10 ± 2°C for seven days, then transferred to a plant growth chamber at 25 ± 2°C for a further seven days under a 12-hour light/dark photoperiod, at which point the final count of emerged seedlings was performed and expressed as a percentage (Cicero and Vieira, 2020).
- Seedling analysis (image analysis): the seeds were set to germinate following the methodology described for the germination test, but without the addition of vermiculite, using four replicates of 20 seeds. Three days after sowing, images were captured and analysed using the GroundEye® system, version S800. Seedlings were carefully removed from the paper and placed in the capture module tray for image acquisition. Following capture, the analysis configuration was set for background colour calibration, with a minimum object evaluation size of 0.08 cm^2^. Image analysis was performed automatically through extraction of mean values for primary root length, based on the methodology of Reis *et al*. (2022).

### Phytotoxicity evaluations

The phytotoxicity index (Pi) of each treatment was calculated relative to the control (seeds without insecticide, of the same genotype and storage period) (Equation 1), adapted from Shahid *et al*. (2024), for the germination variables at 7 days and the cold test (at 4 and 7 days).

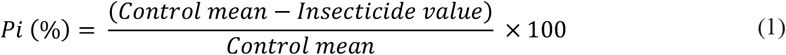

where:

- Control mean: the mean value of the variable analysed for the control treatment at each storage period and genotype.
- Insecticide value: the value of one replicate of the variable analysed for the insecticide treatment, evaluated across the four replicates. Negative Pi (%) values were interpreted as the absence of phytotoxicity and were set to zero for statistical analysis.

### Statistical analysis

- Univariate analysis: data (with the exception of moisture content, for which only means are presented) were first subjected to normality and homoscedasticity tests; when necessary, the most appropriate transformation was selected, followed by analysis of variance (α < 0.05) using the F-test. Where significant, means were compared using the Scott-Knott test (α < 0.05). Analyses were performed using the RStudio ExpDes.pt software package (R Core Team, 2024).
- Multivariate analysis: to provide a clearer visualisation of the interaction among data pertaining to genotypes at storage periods of 0 and 9 months under FD seed treatment, the four replicates of each analysed variable were used (Table 3).

**Table 3.** Treatment codes used in the analysis of mean replicates for each genotype evaluated at 0 and 9 months of storage.

| Genotype | Storage period (months) | Identification (ID) |
| --- | --- | --- |
| H91 | 0 | H91-S0 |
| H44 | 0 | H44-S0 |
| L91 | 0 | L91-S0 |
| L64 | 0 | L64-S0 |
| L44 | 0 | L44-S0 |
| H91 | 9 | H91-S9 |
| H44 | 9 | H44-S9 |
| L91 | 9 | L91-S9 |
| L64 | 9 | L64-S9 |
| L44 | 9 | L44-S9 |

Following treatment coding, the dimensionality reduction technique known as projection pursuit (Friedman and Tukey, 1974) was applied, which identifies patterns through linear projections in lower dimensions from multidimensional data.

Validation of the dimensional reduction is assessed through specification of the objective function (Equation 2), defined as the projection index, a mathematical formulation that expresses the researcher’s interest according to pre-defined objectives, such as cluster visualisation, outlier detection (Equation 3) or treatment discrimination (Equation 4).

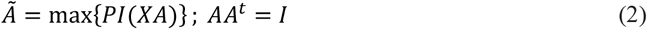

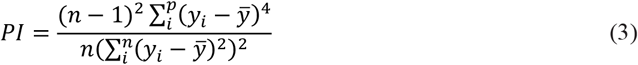

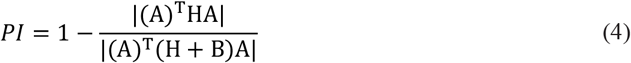

In other words, by specifying a function represented by different PI indices, the technique searches for an orthogonal matrix A that justifies the representation of projections in the d dimension(s) proposed by the index. The projection index used in the present study was the kurtosis index (Equation 3) (Friedman and Tukey, 1974), whose maximisation favours the detection of outliers, where y_i_ (i = 1, …, p) represents the i-th response variable (Ossani *et al*., 2021). Additionally, the index grounded in discriminant analysis theory (Equation 4) was used, which maximises the distance between the means of replicates for each treatment (Peña and Prieto, 2001), where H and B correspond to the between- and within-group covariance matrices of the response variables.

Following these specifications, the MVar.pt package (Ossani and Cirillo, 2025) was used in RStudio, employing the simulated annealing numerical method to optimise the objective function (Equation 2), so as to identify projection spaces in a single dimension that maximise the function for the insecticide treatment groups (FD).

### Scanning electron microscopy (SEM)

Following statistical analyses, the maternal inbred lines L44 and L91 (under both seed treatments) were selected for SEM investigation, evaluating pericarp thickness and the structure of the aleurone layer after 16 hours of imbibition. SEM analysis was performed on a single randomly selected seed from each selected batch, sectioned transversally to the embryonic axis at a standardised region, then mounted on an aluminium stub and coated with gold using a Balzers SCD 050 sputter coater (Silva *et al*., 2017). To examine the samples, the observation region was standardised as the pericarp adjacent to the upper endosperm. The resulting images were used to measure total pericarp thickness (using ImageJ® software) and to assess the integrity of the aleurone layer.

## Results

### Physical analyses

Seed moisture content decreased after two months of storage. However, moisture content was similar amongst genotypes, with variation of less than 2% within the same storage period (Table 4).

**Table 4.**
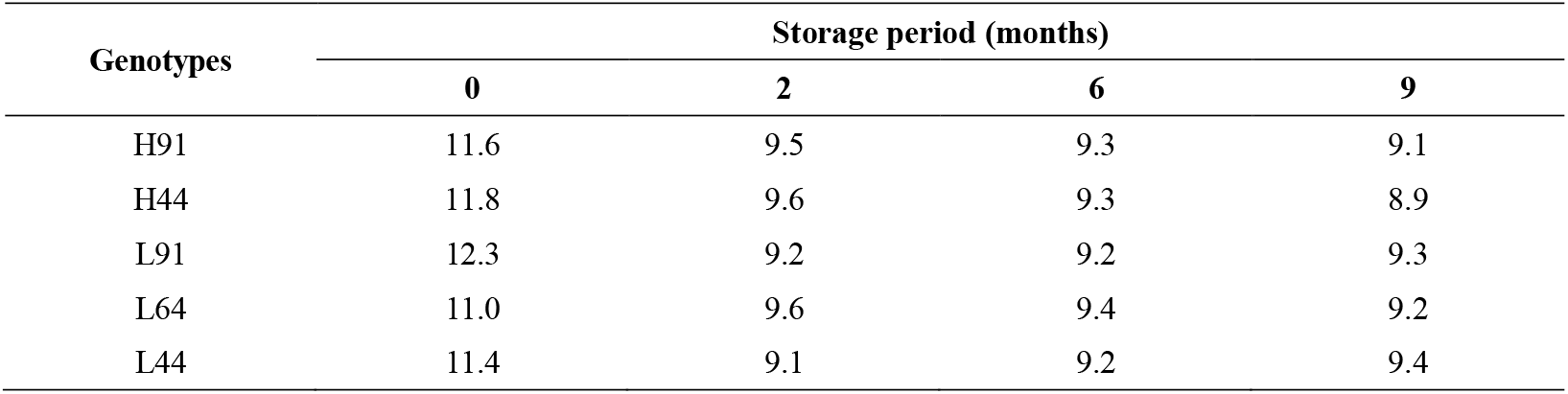
Moisture content (%) of maize seeds of different genotypes during storage, regardless of seed treatment.

### Physiological analyses

Assessment of the first germination count revealed an interaction between genotypes and storage period, as well as an isolated effect of seed treatment (Figure 1).

**Figure 1.**
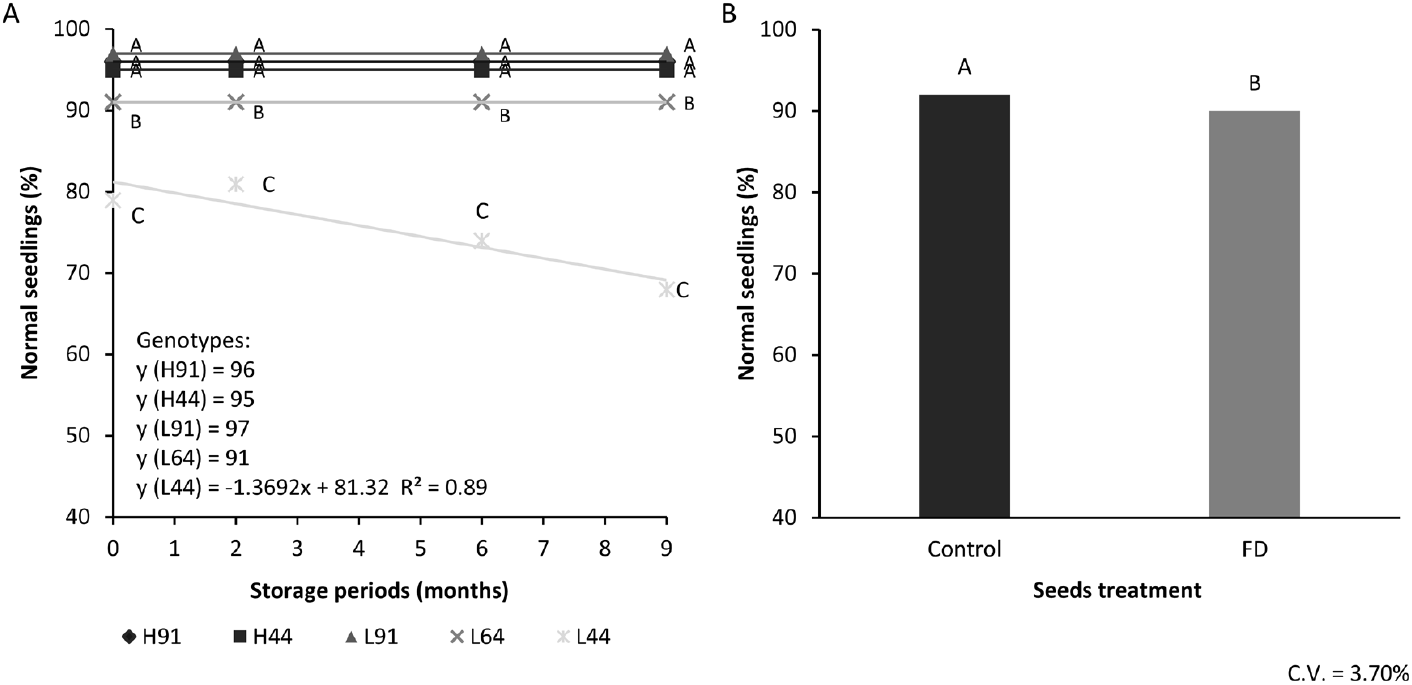
Percentage of normal maize seedlings at 4 days, assessed by the RP+V test: A – effect of genotypes and storage periods; B – influence of seed treatment. *A – Means followed by the same letter within the same storage period do not differ significantly (p > 0.05) by the Scott-Knott test. B – Means followed by the same letter do not differ significantly from each other (p > 0.05) by the F-test.

When genotypes were compared within the same storage period, L44 produced the lowest percentage of normal seedlings, followed by L64. Furthermore, only genotype L44 showed a reduction in the percentage of normal seedlings over the course of storage (Figure 1A). Regarding seed treatments, the insecticide treatment (FD) resulted in a reduction of 2 percentage points (pp) in the percentage of normal seedlings (Figure 1B).

In the assessment of germination at 7 days, a dual interaction was observed between genotypes and seed treatments, as well as an isolated effect of storage period (Figure 2). Genotype L91 was superior under the control seed treatment; however, under the insecticide treatment it performed similarly to the hybrids (H91 and H44). Under both seed treatments, genotype L44 was inferior to all other genotypes; moreover, L44 together with L64 (the male parent) were the only genotypes to show reduced germination following insecticide application, with a reduction of up to 5 pp (Figure 2A). In addition, all genotypes and seed treatments showed a reduction in germination over the storage period, at a rate of approximately 0.5% per 2 months (Figure 2B).

**Figure 2.**
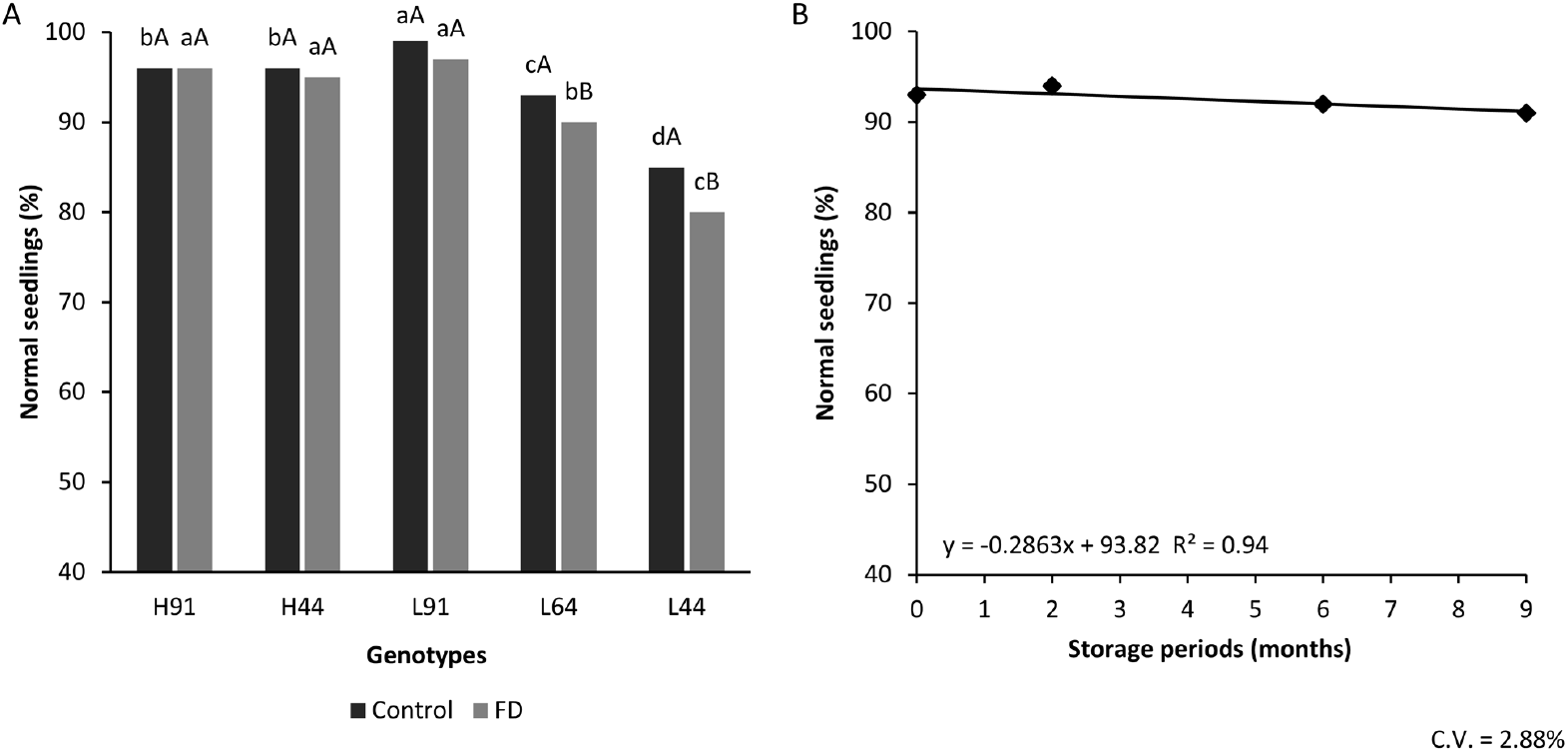
Percentage of normal maize seedlings at 7 days, assessed by the RP+V test: A – effect of genotypes and seed treatment; B – influence of storage period. *A – Means followed by the same letter (lower case among genotypes within the same seed treatment; upper case among seed treatments within the same genotype) do not differ significantly from each other (p > 0.05) by the Scott-Knott test.

Regarding seedling emergence following cold stress, assessed at 4 days after transfer to ideal conditions, a dual interaction was observed between genotypes and storage period, and between genotypes and seed treatment (Figure 3). A similar pattern to that of the first germination count was observed, whereby genotype L44 presented the lowest percentage of normal seedlings over the course of storage, followed by L64. Only genotype L44 showed a significant reduction in the percentage of normal seedlings during storage (Figure 3A).

**Figure 3.**
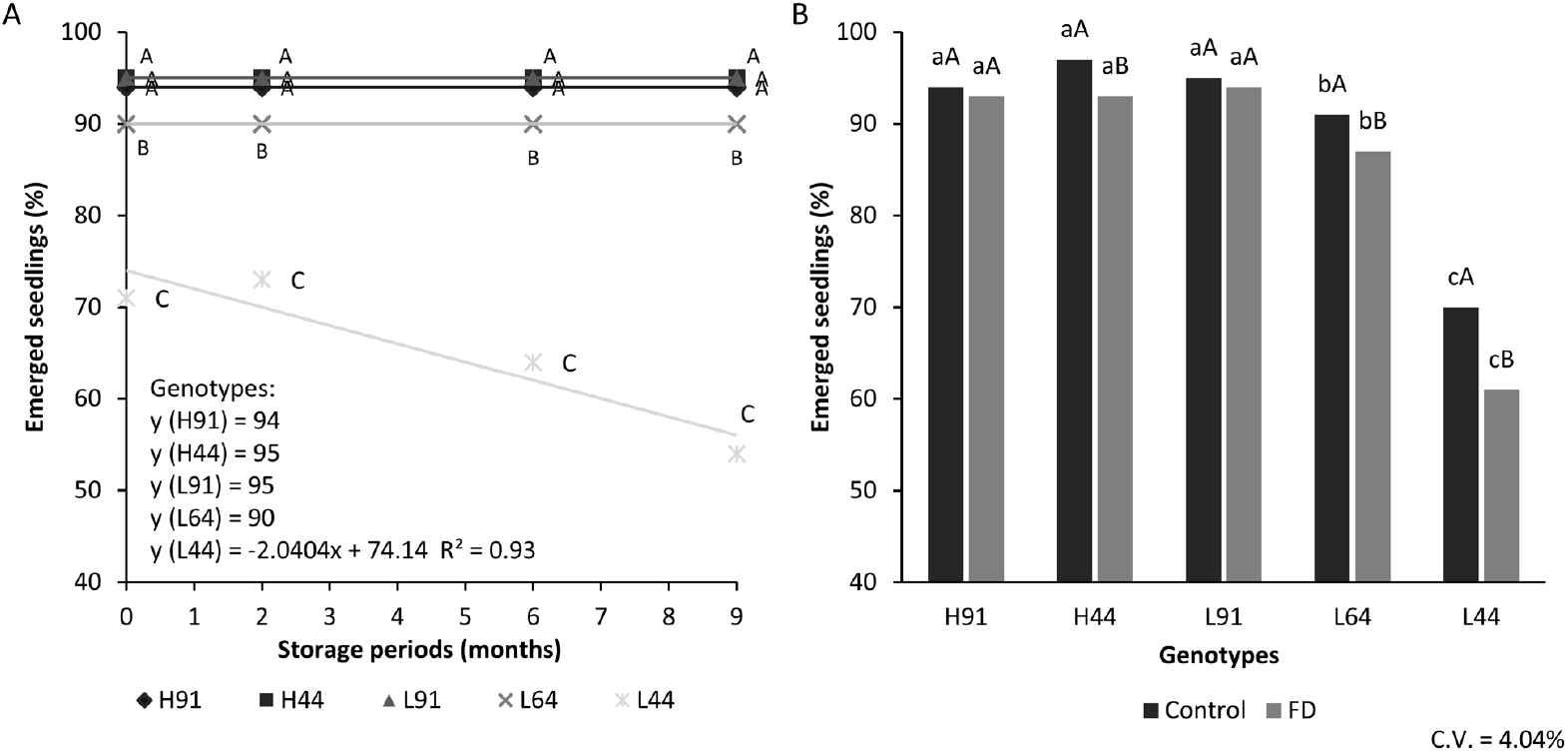
Percentage of maize seedling emergence at 4 days, assessed by the cold test: A – effect of genotypes and storage period; B – effect of genotypes and seed treatment.*A – Means followed by the same letter within the same storage period do not differ significantly from each other (p > 0.05) by the Scott-Knott test. B – Means followed by the same letter (lower case among genotypes within the same seed treatment; upper case among seed treatments within the same genotype) do not differ significantly from each other (p > 0.05) by the Scott-Knott test.

When comparing genotypes across seed treatments, L44 again yielded the lowest results, followed by L64, for both the control and insecticide treatments. However, H44, together with L44 and L64, showed a reduction in the percentage of normal seedlings following insecticide application (Figure 3B).

In the assessment of seedling emergence at 7 days following cold stress, an interaction between genotypes and storage period was observed, as well as an isolated effect of seed treatment (Figure 4). Genotypes L64 and L44 consistently presented the lowest percentages of normal seedlings within the same storage period; however, L44 always exhibited the lowest emergence rate and further showed a reduction in emerged seedlings over the course of storage (Figure 4A). Regarding seed treatments, the FD treatment yielded values 3 pp lower than the control (Figure 4B).

**Figure 4.**
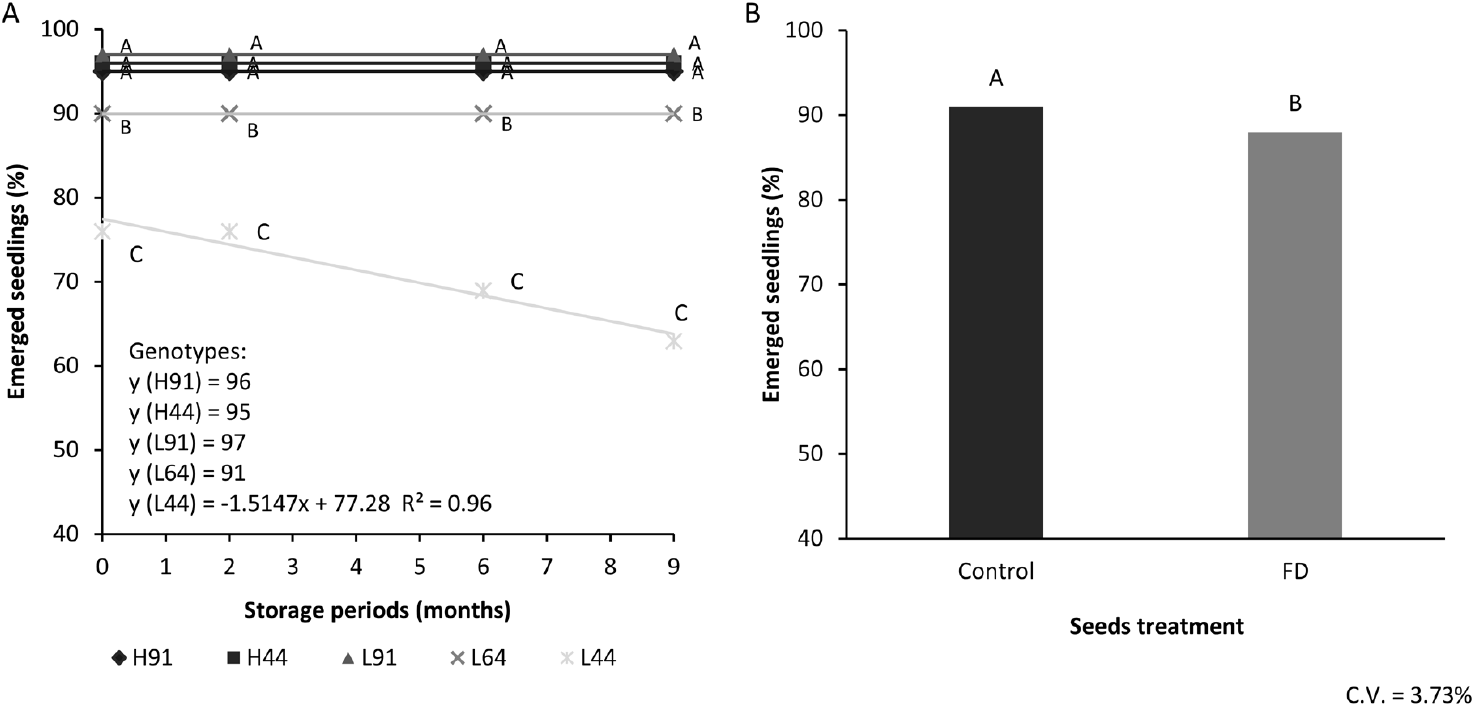
Percentage of maize seedling emergence at 7 days, assessed by the cold test: A – effect of genotypes and storage periods; B – influence of seed treatment.*A – Means followed by the same letter within the same storage period do not differ significantly from each other (p > 0.05) by the Scott-Knott test. B – Means followed by the same letter do not differ significantly from each other (p > 0.05) by the F-test.

With regard to primary root length, genotype L44 under both seed treatments exhibited the lowest root development; the same pattern was observed for genotype L91 when treated with FD (Figure 5A). Regarding storage, primary root length decreased by approximately 0.1 cm per 2 months (Figure 5B).

**Figure 5.**
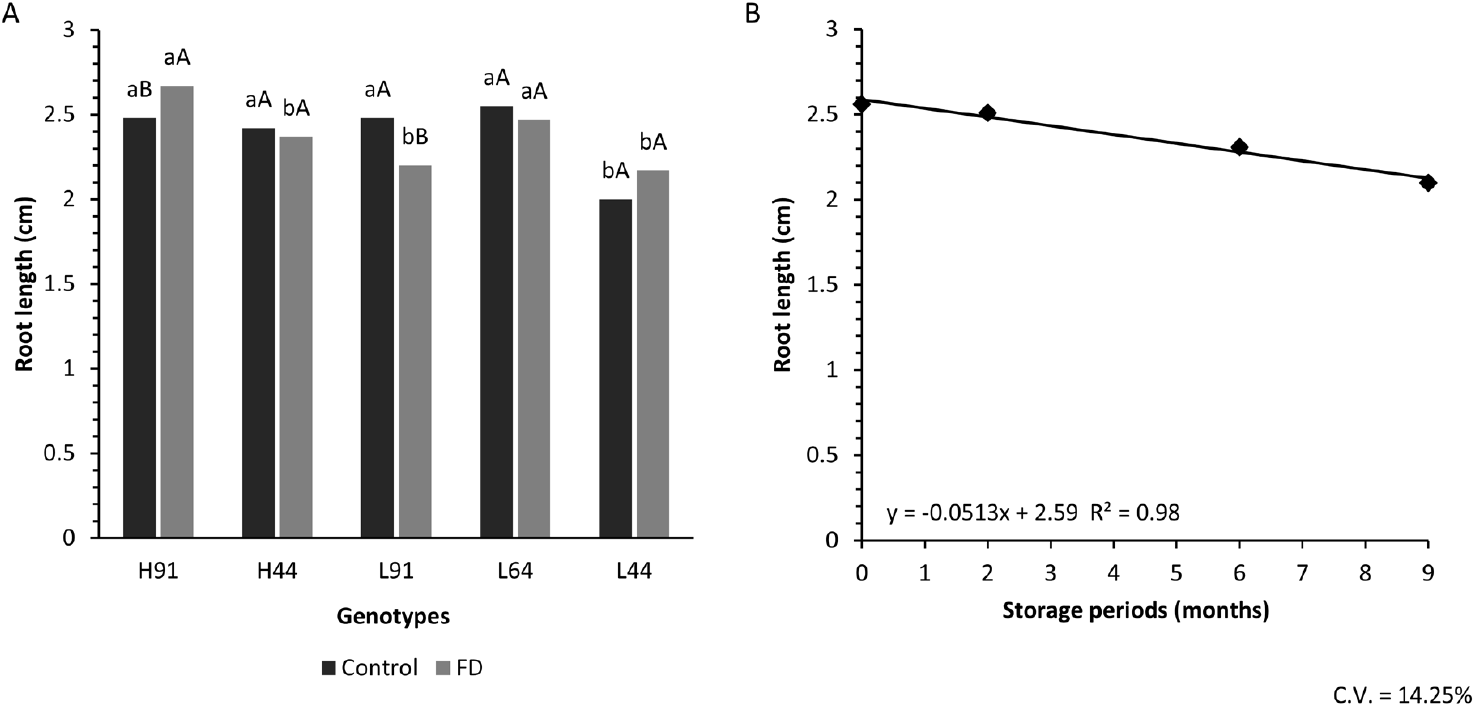
Primary root length (cm) of maize seedlings at 3 days: A – effect of genotypes and seed treatment; B – influence of storage period.*A – Means followed by the same letter (lower case among genotypes within the same seed treatment; upper case among seed treatments within the same genotype) do not differ significantly from each other (p > 0.05) by the Scott-Knott test.

### Phytotoxicity analyses

A significant interaction between genotypes and storage periods was observed for the phytotoxicity index variable (RP+V). In general, genotypes H91, H44 and L91 did not differ from one another. In contrast, genotypes L64 and L44 exhibited the highest phytotoxicity percentages, with L44 being particularly notable, reaching values 15.89 pp and 11.23 pp higher than H91 at 6 and 9 months, respectively. Furthermore, only L64 and L44 showed significant differences across storage periods, evidencing susceptibility to the phytotoxic action of FD over time (Figure 6).

**Figure 6.**
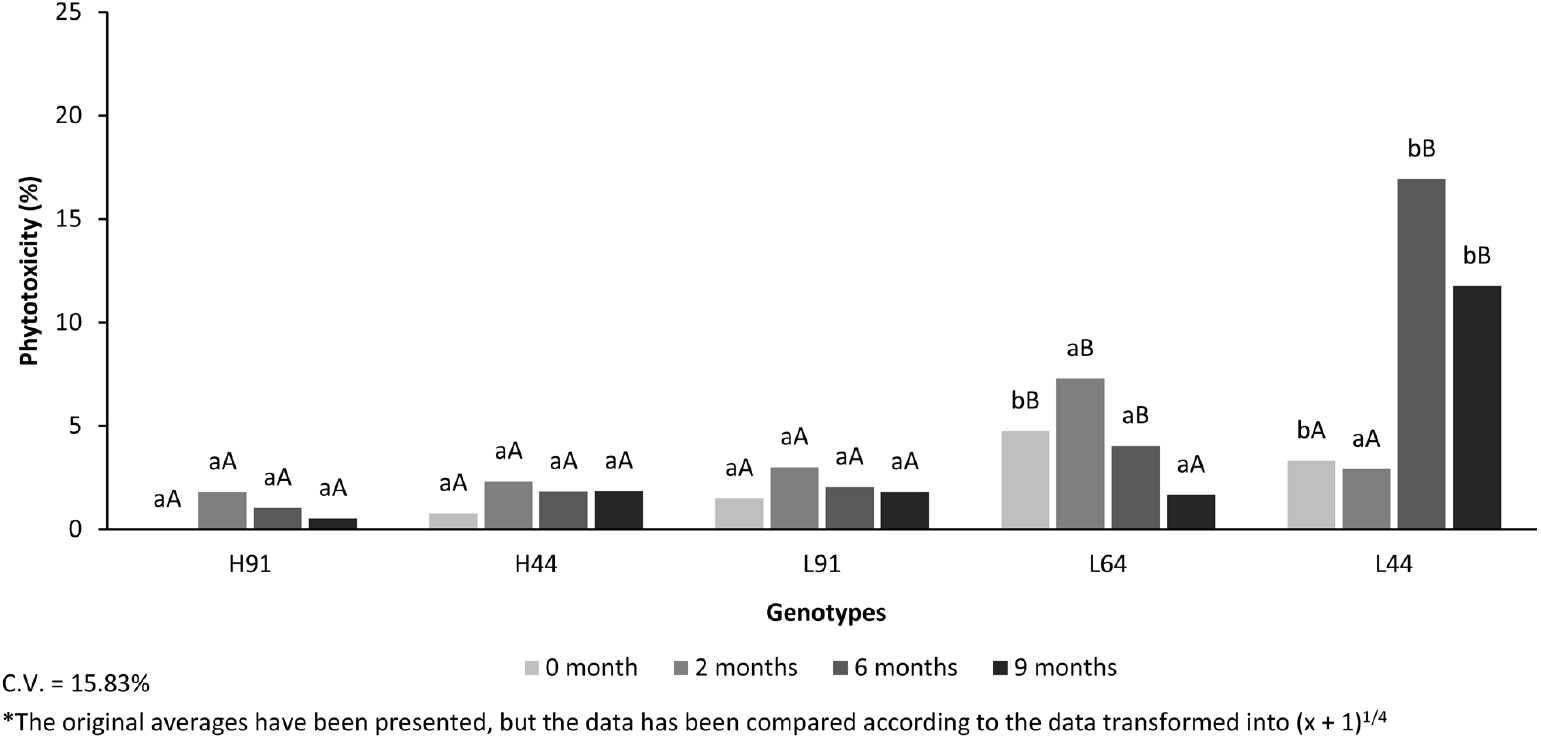
Phytotoxicity (%) in maize seedlings at 7 days, assessed by the RP+V test, effect of genotypes and storage periods. *Means followed by the same letter (lower case among genotypes within the same storage period; upper case among storage periods within the same genotype) do not differ significantly from each other (p > 0.05) by the Scott-Knott test.

Results of the cold test at 4 days revealed significant differences amongst genotypes and storage periods with regard to phytotoxicity (Figure 7). Genotype L44 exhibited the highest phytotoxicity index, with a value 12.64 pp higher than H91. Additionally, H44 together with L64 also showed high phytotoxicity indices (Figure 7A). Regarding storage periods (Figure 7B), the 2-month period stood out for presenting the lowest mean phytotoxicity relative to the other periods.

**Figure 7.**
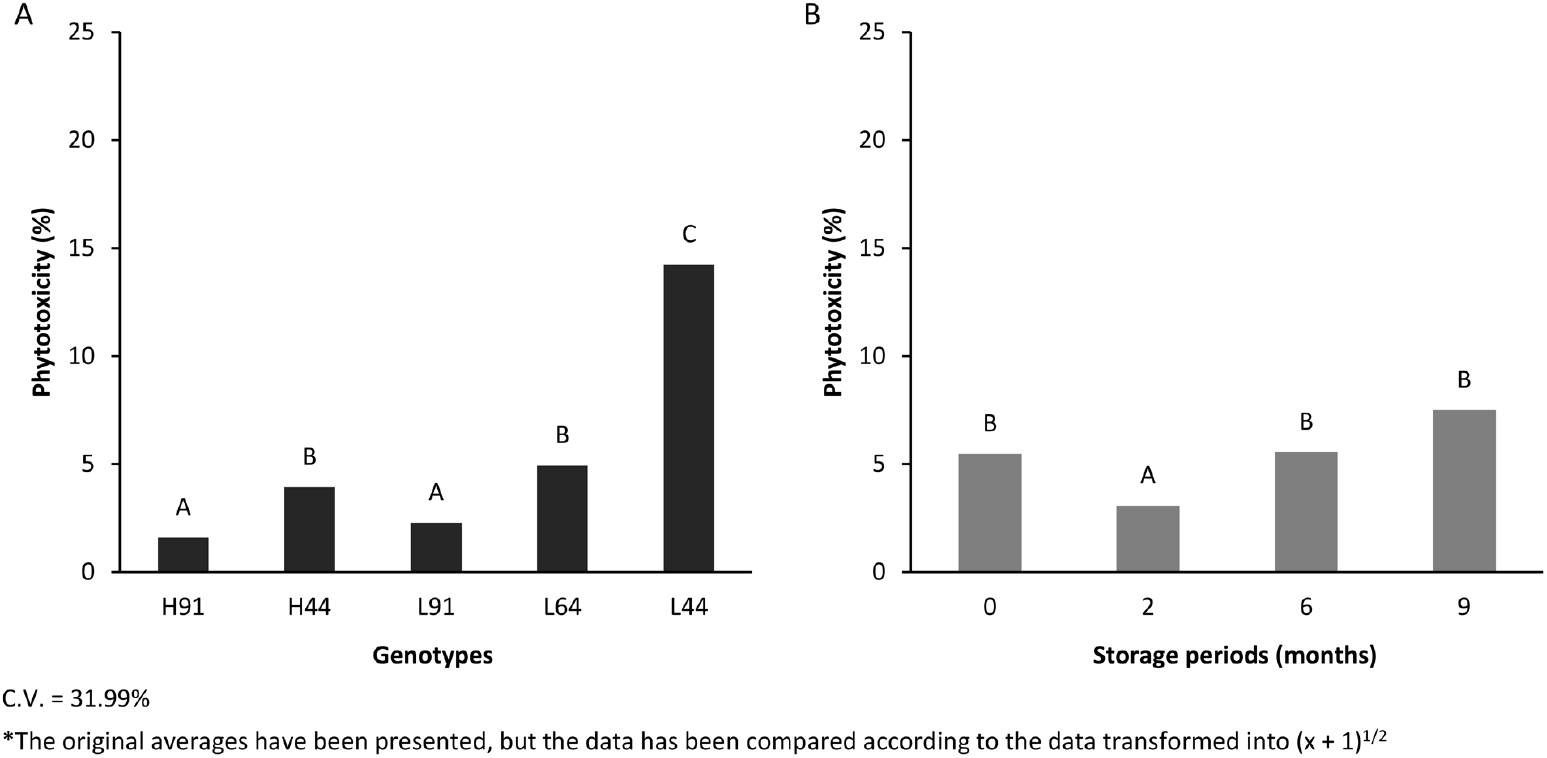
Phytotoxicity (%) in maize seedlings at 4 days, assessed by the cold test: A – influence of genotypes; B – influence of storage periods. *Means followed by the same letter do not differ significantly from each other (p > 0.05) by the Scott-Knott test.

At 7 days of assessment, significant differences were observed amongst genotypes, but not amongst storage periods (Figure 8). With respect to genotypes (Figure 8A), H91, H44 and L91 exhibited the lowest phytotoxicity indices. In contrast, genotypes L64 and L44 showed significantly higher values, 3.85 pp and 6.21 pp, respectively, indicating greater sensitivity to FD relative to H91. No significant difference was observed during storage (Figure 8B).

**Figure 8.**
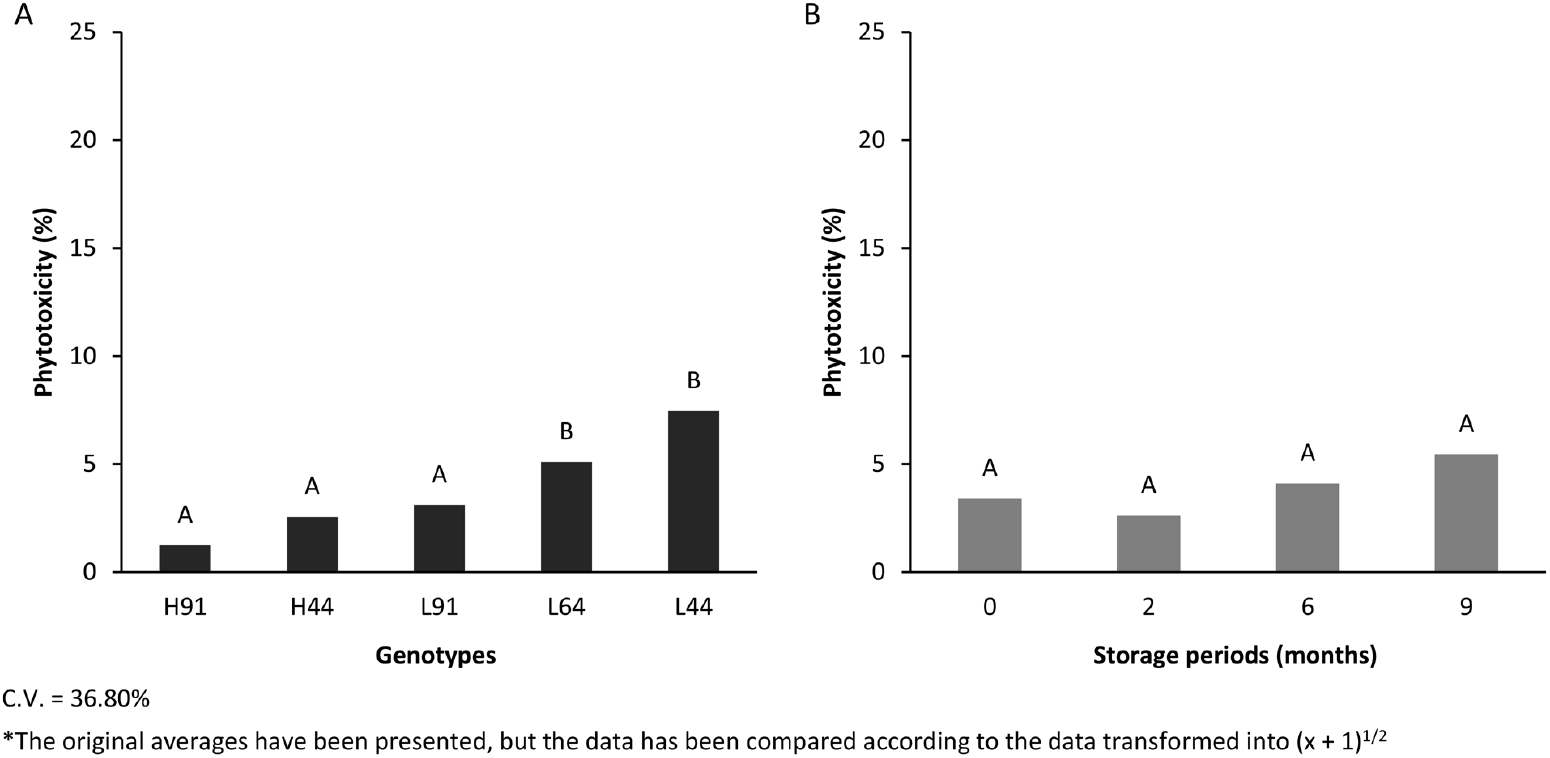
Phytotoxicity (%) in maize seedlings at 7 days, assessed by the cold test: A – influence of genotypes; B – influence of storage periods. *Means followed by the same letter do not differ significantly from each other (p > 0.05) by the Scott-Knott test.

### Multivariate analyses

As illustrated in Figure 9, the projection obtained by maximising the kurtosis index for outlier discrimination, and the treatment discrimination carried out by the index grounded in discriminant analysis theory, which considers the dispersion amongst observations, revealed a prominent highlight for treatment L44-S9, which reflected an extremely high value in the projected space, identifying it as a significant outlier.

**Figure 9.**
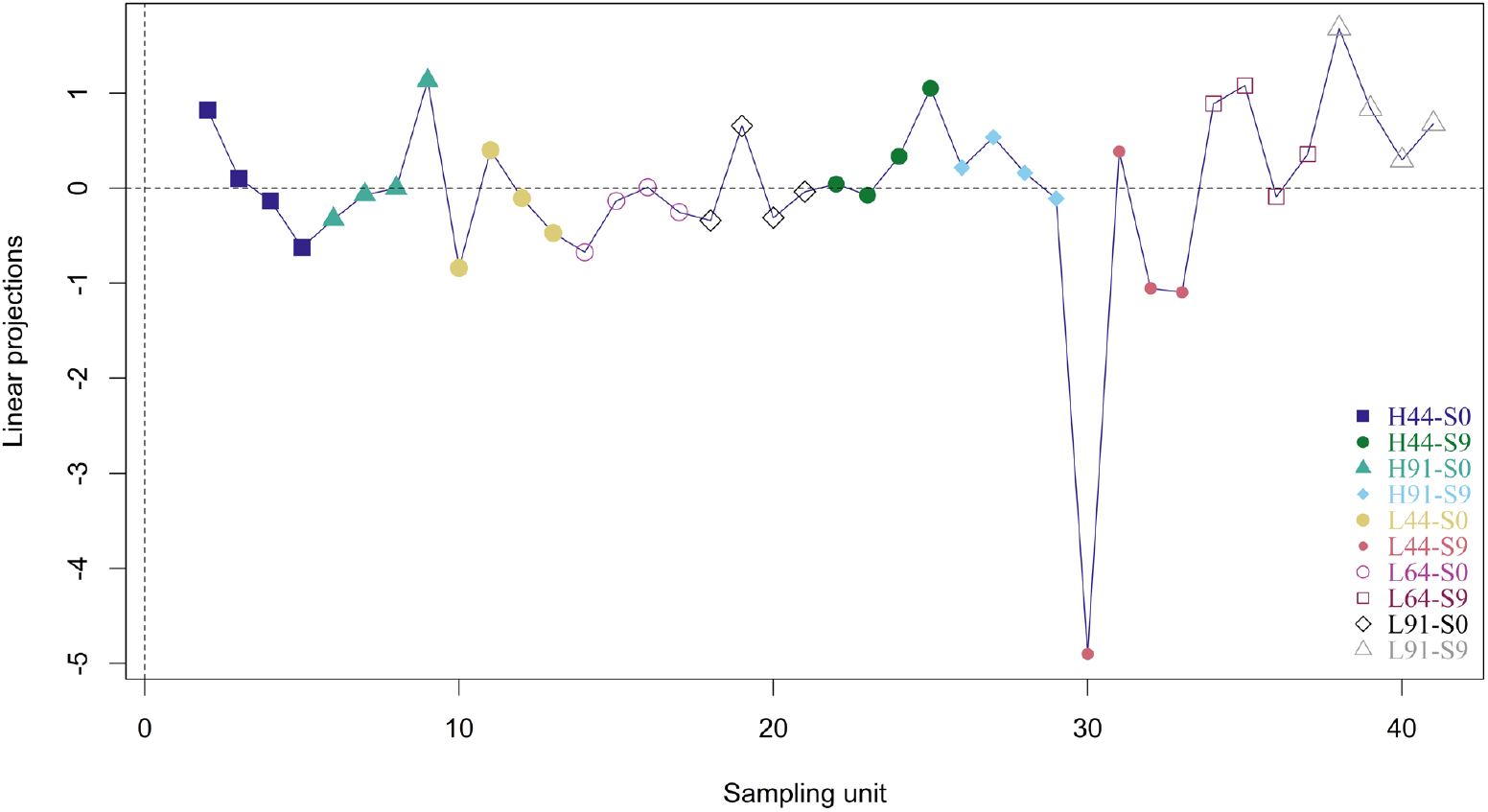
Graph of the projection optimised by the kurtosis index for outlier discrimination.

Linear Discriminant Analysis (LDA) separated the treatments along two main discriminant functions (Figure 10). The first discriminant function (x-axis) was primarily responsible for distinguishing amongst groups. Genotype L44, which exhibited the lowest values across all analysed variables, formed an isolated group at the far right in both storage periods, showing a strong association with the germination vectors (FC_RP+V and RP+V). Conversely, the left side of the plot was occupied by genotypes H91, H44 and L91 (which presented the highest values across the analysed variables), being strongly associated with the phytotoxicity index vectors and CT4.

**Figure 10.**
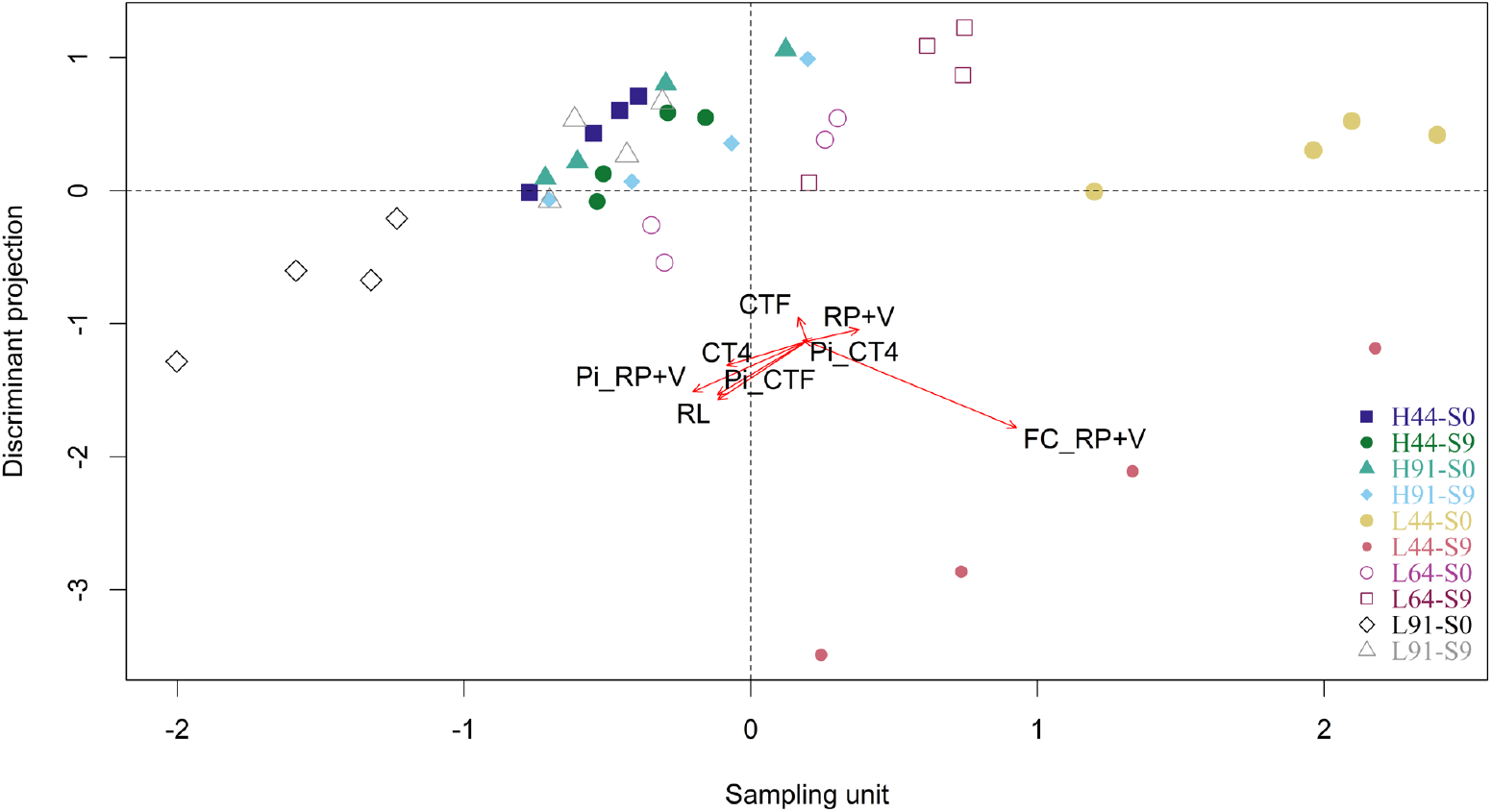
Linear Discriminant Analysis (LDA) of treatments by grouping, based on the analysed variables. *RL = Root length; RP+V = Germination; FC_RP+V = First germination count; CT4 = Cold test (4 days); CTF = Cold test (7 days); Pi_RP+V = Germination phytotoxicity index; Pi_CT4 and Pi_CTF = Phytotoxicity index of the cold test (4 and 7 days).

### Scanning electron microscopy

Scanning electron microscopy (SEM) was used to analyse pericarp thickness in the genotypes following nine months of storage (Figure 11).

**Figure 11.**
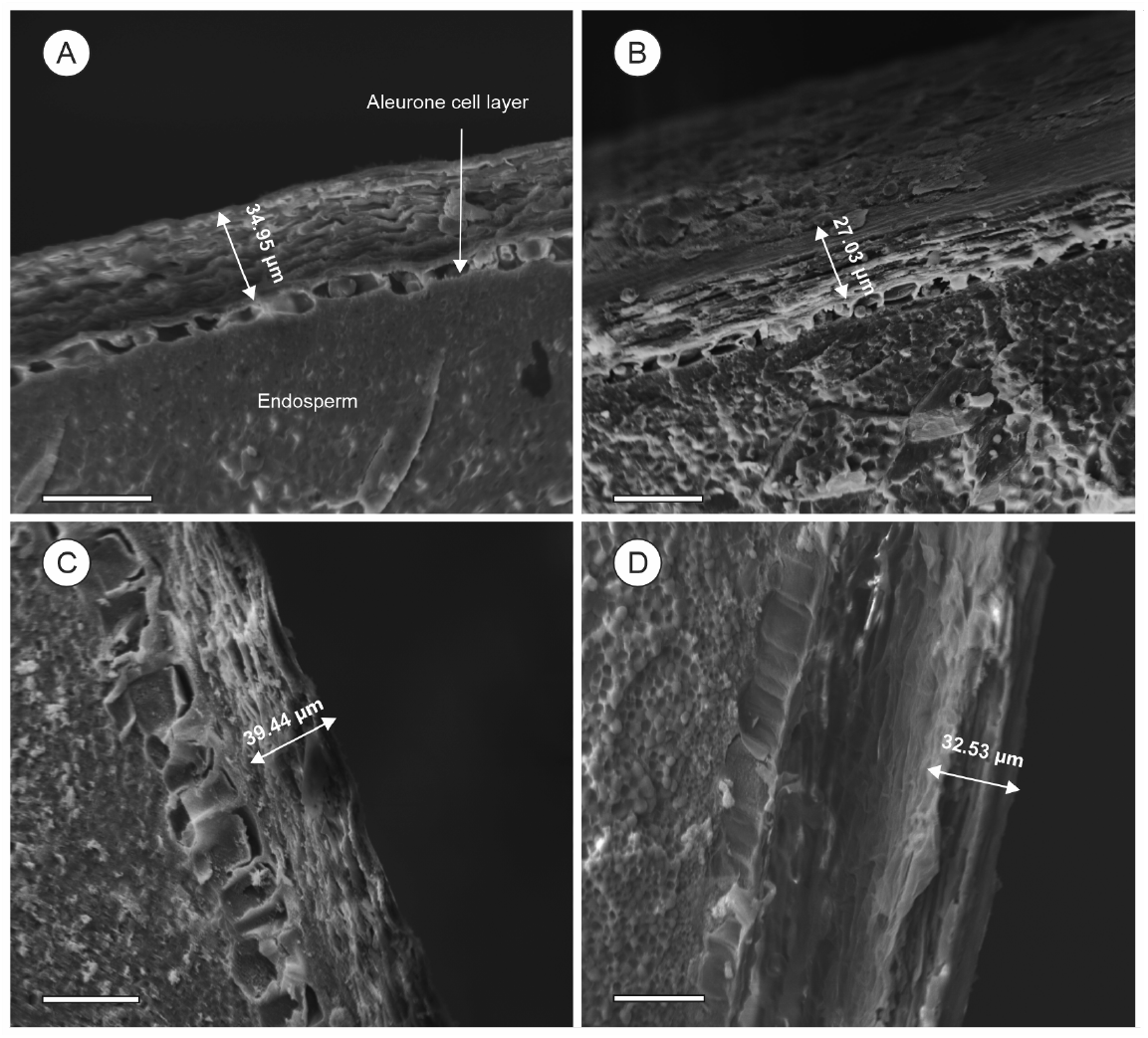
Scanning electron micrograph of pericarp thickness in maize inbred lines under different seed treatments after 9 months of storage. *A: L44-Control; B: L44-FD; C: L91-Control; D: L91-FD. White bar = 20 µm.

Pericarp thickness was reduced under the FD treatment in both genotypes; however, this reduction was more pronounced in L44, which originally presented a thinner pericarp. Genotype L44 under the control treatment (Figure 11A) had a pericarp thickness of 34.95 µm, whilst under the neonicotinoid insecticide treatment (L44-FD, Figure 11B) thickness was reduced to 27.08 µm. For genotype L91, pericarp thickness under the control treatment (Figure 11C) was 39.44 µm, and under the neonicotinoid treatment (L91-FD, Figure 11D) it was reduced to 32.53 µm. These reductions in pericarp thickness, evident in both genotypes, suggest a direct impact of the neonicotinoid insecticide on the structural integrity of the outer pericarp layer following the storage period.

The integrity of the aleurone layer after 16 hours of imbibition was assessed by SEM (Figure 12), providing indicative trends regarding the impact of neonicotinoid insecticide treatment after 9 months of storage.

**Figure 12.**
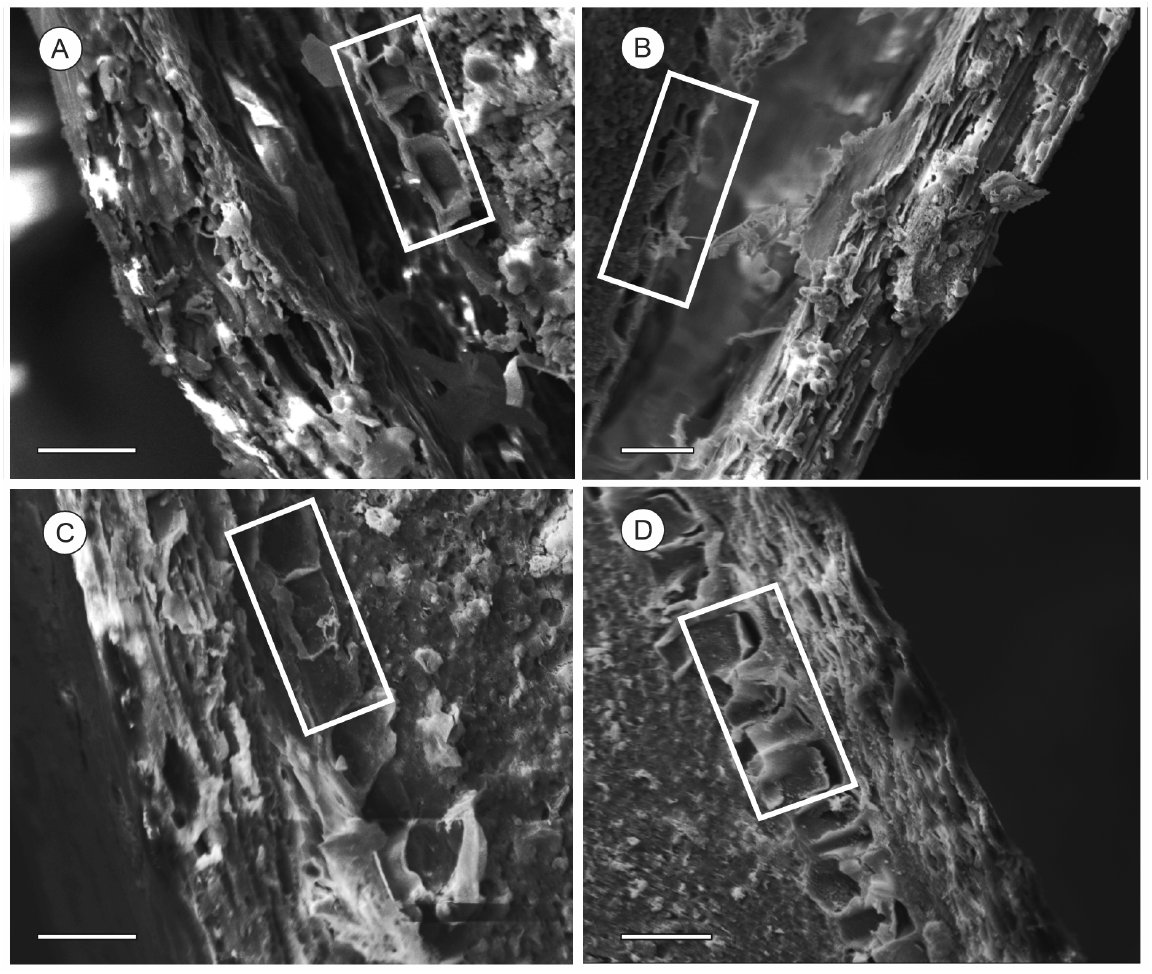
Scanning electron micrograph of aleurone layer cell integrity in maize inbred lines under different seed treatments after 9 months of storage. *A: L44-Control; B: L44-FD; C: L91-Control; D: L91-FD. White bar = 20 µm.

Under the control seed treatment, both L44 (Figure 12A) and L91 (Figure 12C) exhibited a well-preserved aleurone layer with organised cellular structure and cohesive arrangement. In contrast, genotype L44 subjected to the neonicotinoid treatment (L44-FD, Figure 12B) displayed clear signs of disorganisation and potential cellular damage in the aleurone layer, with less cohesive cells and an apparently compromised structure. For genotype L91 under the neonicotinoid treatment (L91-FD, Figure 12D), the aleurone layer maintained a relatively intact organisation, not exhibiting the same degree of structural disruption observed in L44-FD.

## Discussion

Seed treatment is a practice of paramount importance in seed protection, ensuring successful plant establishment in the field (Reis *et al*., 2023). The use of insecticides in ST, particularly those belonging to the neonicotinoid group, has proved to be an effective strategy for the management of early-season pests of the crop, such as *Dalbulus maidis* (DeLong & Wolcott) (Hemiptera: Cicadellidae), which can cause severe yield losses (Neves *et al*., 2022; Machado *et al*., 2025).

On the other hand, neonicotinoids present phytotoxic potential for seeds, reducing their physiological quality, particularly following storage (Oliveira *et al*., 2020; Moraes *et al*., 2022; Reis *et al*., 2026). This behaviour was also verified in the present study, where the use of the FD formulation (which contains a neonicotinoid in its composition) reduced seed quality and consequently increased the phytotoxicity index (Figures 6, 7 and 8), especially after storage.

Furthermore, Tonin *et al*. (2014) and Silva *et al*. (2020) observed that the use of other neonicotinoids, such as clothianidin, imidacloprid and thiamethoxam itself, reduced the quality of maize seeds following storage for 270 days, likely due to their phytotoxic potential. This phytotoxicity also results in alterations to proteins, enzymes and DNA, influenced by the generation of ROS and increased oxidative stress, thereby accelerating the seed deterioration process, as possibly evidenced by the damage caused to the seed pericarp and, subsequently, to embryonic tissues following imbibition, associated with ROS production (Shahid *et al*., 2021).

The reduction in physiological quality was found to be more pronounced following storage; even under the control treatment, a decline in physiological quality was observed, attributable to natural deterioration. Deterioration is commonly described as an irreversible, cumulative and inevitable process (McDonald, 1999), which may lead to the accumulation of cellular damage, resulting in delayed seedling emergence, reduced tolerance to stresses and, ultimately, loss of viability (Zhang *et al*., 2021).

According to Reis *et al*. (2026), limited cellular damage caused by deterioration can generally be repaired, ultimately resulting in a reduction of initial seedling development over the course of storage (Figure 5B). More pronounced deterioration, however, may lead to reduced formation of normal seedlings, whilst extreme deterioration may result in seed death, expressed by a reduction in the percentage of normal seedlings (Figure 2B).

The addition of insecticides to the ST potentiates this deterioration through chemical stress, particularly in susceptible genotypes (L64 and L44). Genotypes L44 and L64 were the only ones to show a significant increase in phytotoxicity, as assessed by the germination test, over the storage period, indicating an inability to mitigate cellular damage over time, with progression towards more severe damage.

Deuner *et al*. (2014) observed that the germination of maize seeds stored under uncontrolled conditions and treated with thiamethoxam was reduced by 16 pp over 12 months of storage; however, only one genotype was evaluated. The present study demonstrates that variability exists amongst genotypes, even when produced under the same conditions, with respect to storage tolerance following ST.

The physiological response of maize seeds to insecticide treatment and subsequent storage is therefore highly genotype-dependent. Genotypes L91, H91 and H44 exhibited high tolerance, maintaining physiological quality, whilst inbred lines L64 and, notably, L44 proved susceptible to phytotoxic effects, particularly during prolonged storage.

Several studies point to genetic constitution as a determinant of high physiological quality and stress response (Marques *et al*., 2019; Costa *et al*., 2021; Melnik *et al*., 2023; Vilela *et al*., 2024). Costa *et al*. (2021) observed that genotypes vary in seed tolerance to delayed drying, indicating that when a susceptible inbred line is used as the female parent, susceptibility is heritable by its progeny, resulting in lower physiological quality.

Furthermore, inbred line L44 exhibited the poorest performance across nearly all physiological and phytotoxicity tests. Whilst univariate analyses revealed differences in specific variables, projection pursuit (multivariate analysis) considered all analysed variables simultaneously and, even so, yielded the same, and even clearer, conclusion: that genotype L44, particularly after 9 months, behaves in a drastically different manner from the remaining genotypes. For genotype L64, differences were detected only in the univariate analyses.

A clear trend observed was the overall greater susceptibility of inbred lines compared with their hybrid offspring. This greater resilience of hybrids is classically attributed to the phenomenon of heterosis, or hybrid vigour (Abreu *et al*., 2019; Wan *et al*., 2022). The combination of different alleles from parental genotypes confers upon hybrids a greater capacity to maintain physiological and metabolic homeostasis under stress conditions, resulting in greater performance stability (Mota *et al*., 2021; Yang *et al*., 2025).

It is important to note that inbred line L91 was an exception, demonstrating remarkable tolerance, often superior to or equivalent to that of the hybrids. This does not invalidate the trend but rather enriches it, showing that high tolerance to post-treatment storage is not exclusive to hybrids. The performance of L91 may be associated with specific tolerance genes that, when present in homozygosity in this line, are highly effective, such as the *ZmDREB2A/2*.*1S* gene, related to drought tolerance characteristics (Vilela *et al*., 2024).

The overall superiority of hybrids reinforces their commercial importance by ensuring greater reliability and field performance. However, the existence of a highly tolerant parental line such as L91 is of considerable value for plant breeding, as it may serve as a proven source of tolerance genes for the development of future hybrids with even greater storage tolerance following ST.

Moreover, the tolerant line L91 naturally possesses a thicker pericarp (39.44 µm) compared with the susceptible line L44 (34.95 µm). Insecticide treatment reduced pericarp thickness in both genotypes, but the effect was more pronounced in L44. The maize seed pericarp exhibits selective permeability, allowing the passage of non-ionic compounds (such as neonicotinoids) whilst restricting ionic ones (Diaz *et al*., 2014). A thinner pericarp may therefore permit faster and/or greater penetration of the active ingredient, leading to a reduction in physiological quality, as observed in the case of L44.

In addition, the aleurone layer cells of L44 treated with neonicotinoids exhibited disorganisation and disruption, whilst those of L91 remained relatively intact and organised (Figure 12). This may be explained by the fact that increased chemical stress from insecticides leads to ROS accumulation, which promotes deterioration, evidenced by cellular collapse, mitochondrial membrane deconstruction, elevated genetic material damage and high ROS accumulation, ultimately resulting in a reduction of physiological quality (Ebone *et al*., 2019; Reis *et al*., 2026)

Furthermore, although inbred line L64 was the male parent of both hybrids, hybrid H44, derived from the susceptible line L44 as the female parent, exhibited some sensitivity to the insecticide in the cold test at 4 days. This behaviour may be related to the genetic contribution of L44, which potentially transmitted susceptibility genes. According to Zhang *et al*. (2023), genes found in the pericarp are predominantly of maternal origin (97%), which may account for the results observed for genotype H44 treated with FD in the cold test at 4 days following storage (Figures 3B and 7A).

Pericarp thickness may therefore potentially serve as an effective marker of storage tolerance following treatment in breeding programmes; however, further studies are still required.

## Conclusions

Tolerance of maize seeds to neonicotinoid seed treatment during storage varies significantly amongst genotypes, being more pronounced in inbred lines than in hybrids.

Inbred lines L44 and L64 were the most susceptible genotypes, whilst L91, H91 and H44 maintained physiological quality throughout storage.

Pericarp thickness is a potential morphological marker for selecting maize genotypes tolerant to neonicotinoid seed treatment in breeding programmes.

## Author contributions

V.U.V.R. conceived and designed the research, conducted the experiments, analysed the data and wrote the paper. G.I.S.T. and M.S.R.P. assisted in data collection. S.A.G.A. and M.A.C. assisted in statistical analysis and manuscript revision. G.A.S. performed the scanning electron microscopy analysis and contributed to manuscript revision. E.R.C. supervised the research, reviewed the project and provided administrative and financial support. All authors have read and agreed to the published version of the manuscript.

## Acknowledgements

The authors are grateful to the CNPq, FAPEMIG, and the CAPES for financial support, scholarships and research productivity grants. The authors also thank the Laboratory of Electron Microscopy and Ultrastructural Analysis of the Federal University of Lavras, and Finep, for supplying equipment and technical support for the scanning electron microscopy experiments. Special thanks to Seedcare Institute Syngenta Brazil for technical support.

